# EPySeg: a coding-free solution for automated segmentation of epithelia using deep learning

**DOI:** 10.1101/2020.06.30.179507

**Authors:** Benoit Aigouy, Benjamin Prud’Homme

**Affiliations:** Aix Marseille Univ, CNRS, IBDM, Marseille, France

## Abstract

Epithelia are dynamic tissues that self-remodel during their development. At morphogenesis, the tissue-scale organization of epithelia is obtained through a sum of individual contributions of the cells constituting the tissue. Therefore, understanding any morphogenetic event first requires a thorough segmentation of its constituent cells. This task, however, usually implies extensive manual correction, even with semi-automated tools. Here we present EPySeg, an open source, coding-free software that uses deep learning to segment epithelial tissues automatically and very efficiently. EPySeg, which comes with a straightforward graphical user interface, can be used as a python package on a local computer, or on the cloud via Google Colab for users not equipped with deep-learning compatible hardware. By alleviating human input in image segmentation, EPySeg accelerates and improves the characterization of epithelial tissues for all developmental biologists.

## Introduction

Epithelia are dynamic tissues undergoing dramatic shape changes throughout their development. A prerequisite for understanding these morphogenetic events is the thorough segmentation of cells constituting the tissue. To this aim, numerous semi-automated methods have been developed ^1–4^ but they require time-consuming manual correction to achieve optimal segmentation.

Over the past few years, deep learning and more particularly convolutional neural networks (CNNs) revamped the computer vision field, including image segmentation by alleviating the need for user correction of the segmentation. The advent of simple programming frameworks, such as Keras and TensorFlow^5,6^, made deep learning accessible to most developers but still excludes people lacking coding skills, preventing deep learning from being broadly adopted by the scientific community. Few attempts to bring convolutional neural networks to well-known image processing frameworks such as ImageJ or FIJI exist ^7–11^, but they require an up-to-date and adequately configured computer. More importantly, most often those powerful yet very poorly generalizable CNNs need to be trained *de novo* on user data to work efficiently. Unfortunately, such training cannot be done directly in FIJI/ImageJ and requires, again, coding. So far, little effort has been made to facilitate CNN training and use by regular users ^12^.

To address all these limitations, we present EPySeg, a coding-free solution to segment raw images of epithelial tissues very efficiently, using a set of pre-trained generalist neural networks. Also, EPySeg comes with a complete and straightforward graphical user interface (GUI), allowing for users curious about deep learning as well as more advanced ones to build and train custom networks to achieve any desired segmentation paradigm. EPySeg is available at https://github.com/baigouy/EPySeg and a minimal version can also be used on Google Colab (https://github.com/baigouy/notebooks) for users equipped with low-end graphic cards.

## Results

In this study, we set to develop a software that uses deep learning to automate the time-consuming segmentation of 2D epithelial tissue images. We selected Linknet architectures because they are known to perform well at image segmentation tasks ^13^ **(see also material and methods).** Our networks were trained on large amounts (i.e. big data) of high-quality human segmented images. Practically, the network aims at reproducing such high-quality segmentation by minimizing a loss function. To do so, the network must learn characteristic features from the user-provided input, and use this knowledge to output a valid segmentation **(Fig. 1).** The segmented epithelia used to train our networks were generated with the watershed algorithm ^14^ and manually edited to remove segmentation artefacts **(see material and methods).** Since human input is limited solely to editing, segmentation variability is low and should improve network training when compared to purely handmade segmentation. Also, in order to allow the networks to segment virtually any 2D epithelium, we trained them on images of highly divergent epithelia acquired using several microscopy set-ups **(see material and methods).**

**Figure 1:**
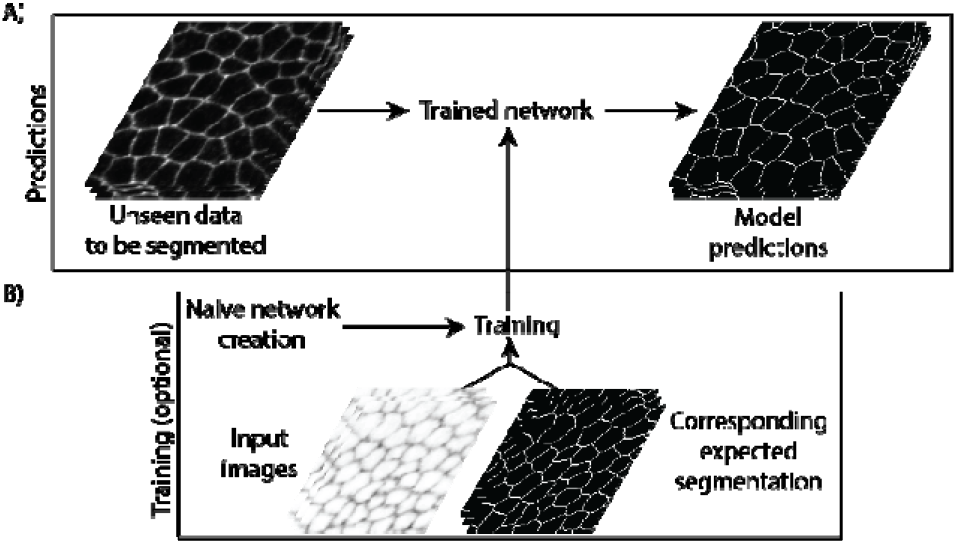
Deep learning segmentation pipeline A) Unseen images of epithelia are provided to a trained neural network that returns a segmentation prediction. B) Convolutional neural networks can be built from scratch then trained (optional). Training requires original images and their expected segmentation to be fed to the network. Note that both predictions and training can be done on the local computer using the EPySeg GUI or the cloud using Google colab.

EPySeg efficiently segments 2D epithelial cells from different tissues imaged with different optics **(Fig. 2 and table 1).** On average, it outperforms existing tools such as Cellpose ^15^ on most epithelia in two ways:, its approximation of the cell outline is more precise than that of Cellpose, and it also loses fewer cells **(Fig. 2).** We note, however, that unlike Cellpose, EPySeg is not able to segment cells in culture **(Table 1)** since it was not trained to accomplish such task.

**Figure 2:**
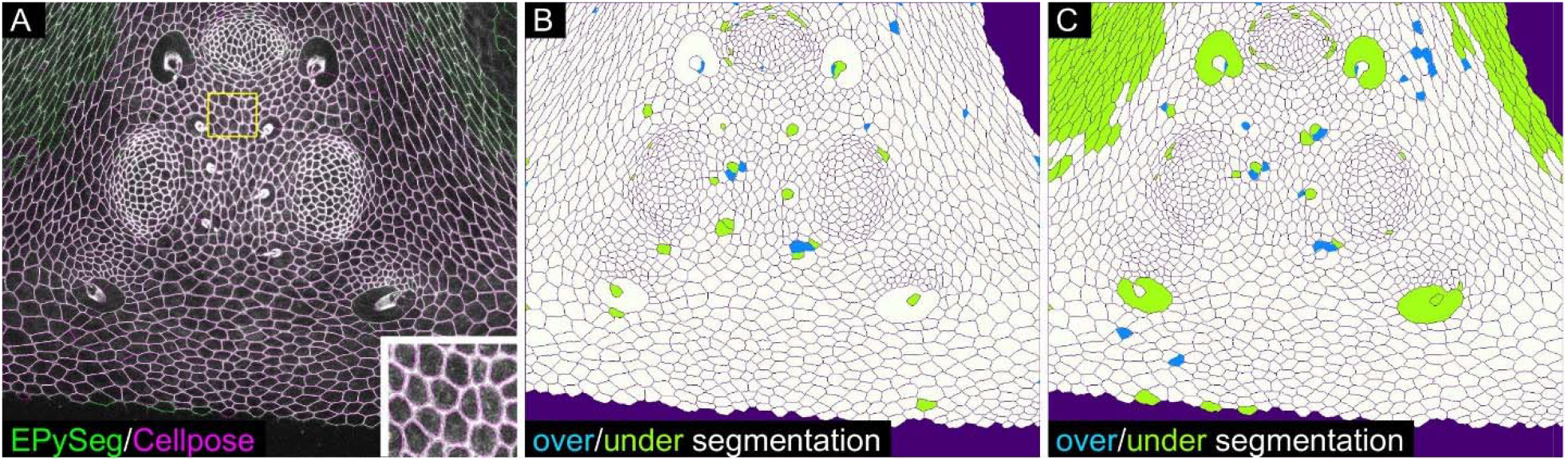
EPySeg vs. Cellpose segmentation of an unseen epithelium image A) Segmentation of the image of the fly ocelli, an unseen epithelium, by EPySeg (green) and cellpose (purple) overlaid over the original input image. The inset shows a magnified view corresponding to the ROI shown in A; note that cellpose cell outlines lie at a distance from the real cell boundaries. Comparison of B) Epyseg and C) Cellpose over (blue) and under segmentation (green) to the ground truth segmentation (see also table 1 for additional quantifications of unseen samples). Correct segmentation is shown in white.

Finally, to make our epithelial segmentation tool easily accessible to a broad audience, we created a graphical user interface (GUI) and a documentation for it (https://github.com/baigouy/EPySeg). This interface allows for building, training and running convolutional neural networks. It is built in a way that non-experts can rely on the default settings to get a decently trained network and gain knowledge about deep learning, while the advanced users can visually fine-tune parameters to get optimal results. Also, since the majority of scientific computers are not deep learning-ready, making it difficult to train convolutional neural networks, we provide a minimal user interface to run EPySeg on Google Colab, along with online guidelines, to allow for everyone to use it (https://github.com/baigouy/notebooks).

## Material and methods/supplement

### Recommended equipment

Convolutional neural networks were trained on a 64 GB RAM Dell precision 7820 equipped with a Nvidia GeForce® RTX 2070 graphic card with 8 GB RAM. Most training lasted less than 12 hours. We flawlessly trained and ran our pre-trained networks on Google Colab, hereby providing a good alternative for users with deep learning-incompatible systems.

### Data

Convolutional neural networks were trained on several *Drosophila* epithelia stained with E-cadherin-GFP that diverged largely from one another. At embryonic stage E-cad staining appeared dotted ^16–18^ and the boundary to cytoplasm signal ratio was low. During pupal wing development staining appeared continuous and presented a higher boundary to cytoplasm ratio except for stretched cells. Finally, our third training set contained images of the fly abdomen including giant, polyploid, larval cells larval and tiny histoblast nest cells ^19^, in order to have a network that segments cells without size bias. Input images were max or stack focuser projections ^20^ of confocal stacks of epithelial tissues. Segmented cell outlines serving as ground truth for training the network, were generated using the watershed algorithm of Tissue Analyzer ^1,14^. Two of the datasets were acquired on regular Leica or Zeiss confocal microscopes **(Leica SP5 and LSM 510),** while the third training set was acquired on a spinning disc **(Roper)** to further increase the variability between images. Importantly, we paid a lot of attention to the quality of the segmentation masks fed to the convolutional neural network and cropped out regions where segmentation quality was poor as well as regions that were not segmented (e.g. cells adjacent to the tissue of interest) in order not to perturb the learning process.

### Data augmentation

Given the relatively small size of our training set for deep learning (images /cells) and to prevent the neural network from overfitting, we used data augmentation. Practically, we randomly applied the same deformation (rotation, translation, shear, magnification, flip, …) to the input and output images. Our data augmentation algorithm currently supports 2D and 3D images (only 2D images were used in this study).

### Convolutional neural network building and training

Convolutional neural networks rely on the TensorFlow and Keras tools and were generated using the segmentation_models library from **(https://github.com/qubvel/segmentation_models).** Typically, we used Linknet^13^ architectures and varied the encoder layers. We found the vgg16 and seresnext101 encoders ^21,22^, both known to perform well at classification tasks, very efficiently segment epithelia. Networks were trained for 100 to 300 epochs on the complete train set at every epoch. Depending on their memory requirements, networks were trained with a batch size varying between 2 and 64 images and with a tile size ranging from 64 to 256 pixels in width and height. We chose the intersection over union (IoU) also called the Jaccard index for the loss function as it is particularly well suited to evaluate differences between binary images.

### Segmentation quantifications

As a measure for the overlap between neural network-generated cell outlines and user-provided ones, we computed the IoU score. The IoU measures ranges between 0 and 1, with 1 being identical images and 0 being images having nothing in common. Of note, a value of 0.5 for the IoU score is generally considered as being a good match between two binary images. Importantly, a poor match between cell outlines is not necessarily an indicator of poor image segmentation, indeed, one can imagine a case where the central part of the cell is always properly detected while the edges are ill-defined. To get an estimate of the quality of the segmentation and to evaluate over- and undersegmentation in an image, we designed a second metric. We define this so-called ‘segmentation quality’ measure, as the count of properly segmented cells minus the number of over- and under-segmented cells, and divide the result by the total number of cells in the human-corrected image. We define properly segmented cells as cells having an IoU score superior to 0.7 when comparing the cytoplasm generated by the neural network to the user one. The segmentation quality measure has a maximum value of 1, which is reached when all cells are properly segmented without over- or under-segmentation. This measure can become close to 0 or even negative, this indicates that lots of cells have not been properly identified and that tremendous human editing is required. For optimal comparison to Cellpose, we let the Cellpose software compute the optimal cell size before segmenting the tissues.

### Software

The software was entirely coded in Python 3. The graphical user interface was made with PyQT5 (Riverbank®). The source code of our tool along with install instructions can be found at the following link (https://github.com/baigouy/EPySeg).

